# Cross-Modality and Self-Supervised Protein Embedding for Compound–Protein Affinity and Contact Prediction

**DOI:** 10.1101/2022.07.18.500559

**Authors:** Yuning You, Yang Shen

## Abstract

**Motivation:** Computational methods for compound–protein affinity and contact (CPAC) prediction aim at facilitating rational drug discovery by simultaneous prediction of the strength and the pattern of compound–protein interactions. Although the desired outputs are highly structure-dependent, the lack of protein structures often makes structure-free methods rely on protein sequence inputs alone. The scarcity of compound–protein pairs with affinity and contact labels further limits the accuracy and the generalizability of CPAC models.

**Results:** To overcome the aforementioned challenges of structure naivety and labelled-data scarcity, we introduce cross-modality and self-supervised learning, respectively, for structure-aware and task-relevant protein embedding. Specifically, protein data are available in both modalities of 1D amino-acid sequences and predicted 2D contact maps, that are separately embedded with recurrent and graph neural networks, respectively, as well as jointly embedded with two cross-modality schemes. Furthermore, both protein modalities are pretrained under various self-supervised learning strategies, by leveraging massive amount of unlabelled protein data. Our results indicate that individual protein modalities differ in their strengths of predicting affinities or contacts. Proper cross-modality protein embedding combined with self-supervised learning improves model generalizability when predicting both affinities and contacts for unseen proteins.

**Availability:** Data and source codes are available at https://github.com/Shen-Lab/CPAC.

**Supplementary information:** Supplementary data are included.

## 1 Introduction

Most FDA-approved drug–target pairs are between small-molecule compounds and proteins (Santos *et al.*, 2017). Considering the enormous chemical space that is estimated to contain 10^60^ “drug-like” compounds (Bohacek *et al.*, 1996), it is desirable to virtually screen compounds with high throughput and high accuracy, based on their computationally predicted properties as well as interactions with proteins (off)targets. Thanks to quickly growing data, modeling techniques, and computing power, many machine-learning and deep-learning methods emerge for predicting compound–protein interactions, in particular, the structure-free ones addressing the often unavailability of protein structures (Öztürk *et al.*, 2018; Gao *et al.*, 2018; Karimi *et al.*, 2019, 2020; Tsubaki *et al.*, 2019; Jiang *et al.*, 2020; Li *et al.*, 2020).

Recent progress in structure-free methods includes increasing resolution of what they predict: from binary interactions (Gao *et al.*, 2018; Tsubaki *et al.*, 2019) to continuous affinity or activity values (Öztürk *et al.*, 2018; Karimi *et al.*, 2019). The progress also includes increasing explainability about how they predict such interactions: intermolecular atom–residue non-bonded contacts underlying compound– protein affinities are additionally predicted, often by introducing (Gao *et al.*, 2018; Karimi *et al.*, 2019), regularizing (Karimi *et al.*, 2020), and supervising (Karimi *et al.*, 2020; Li *et al.*, 2020) various attention mechanisms. We refer to such an explainable affinity prediction problem as compound–protein affinity and contact (CPAC) prediction.

Despite the aforementioned progress, two challenges present major barriers to the accuracy and the generalizability. (**i**) **Lack of structure awareness.** While being generally applicable by assuming no co-crystal, docked or even unbound structures as protein inputs, structure-free methods rely on 1D amino-acid sequences (Öztürk *et al.*, 2018; Li *et al.*, 2020) and sequence-predicted 1D structural property sequences (Karimi *et al.*, 2019), thus lack the awareness of 3D structures that are critical to what they predict (affinity and contact labels). (**ii**) **Scarcity of labelled data.** Compared to the daunting size of compound–protein pairs, only a tiny fraction are labelled with affinity measurements and even less are labelled with non-bonded atomic contacts from co-crystal structures. This challenge for supervised models is known as “supervision starvation”.

To address the aforementioned challenges, we make two major contributions accordingly. First, to address structure naivety, without demanding co-crystal, compound-docked, or even unbound protein 3D structures, we consider protein data as available in both modalities of 1D sequences and sequence-predicted 2D graphs (contact maps). Recent revolution in protein structure prediction (Jumper *et al.*, 2021; Baek *et al.*, 2021) is making the structure modality increasingly available. We introduce various neural network architectures to separately or jointly embed protein modalities and introduce **cross-modality learning** to inject structure-awareness into resulting protein embeddings. Two cross-modality strategies, concatenation and cross interactions, are introduced to encode the modalities independently and dependently. Second, to address supervision starvation, without demanding more labelled data, we leverage massive unlabelled protein data and introduce various **self-supervised learning** strategies to pre-train protein embedding. Specifically, we use masked language models (Devlin *et al.*, 2018) for pre-training protein sequence embedding and graph completion and graph contrastive learning (You *et al.*, 2021) for pre-training protein contact-map embeddings.

In cross-modality learning, we ask whether individual modalities could excel in predicting either affinities or contacts as well as whether and how their individual strengths could be combined for better accuracy and generalizability. Our results indicate that the 1D and 2D modalities of protein data do not dominate each other in CPAC prediction for proteins seen in the training set; however, they tend to generalize better for unseen proteins in affinity prediction and contact prediction, respectively. We thus provide a conjecture for such observations, which is verified numerically. To integrate knowledge from 1D and 2D protein modalities, two cross-modality schemes are proposed, with empirical demonstration that they achieve the state-of-the-art (SOTA) performance.

In self-supervised learning we ask how to design self-supervised strategies, within and across individual protein modalities, in order to improve model accuracy and generalizability. We leverage rich unlabelled protein data and adopt self-supervised techniques for sequences and graphs so as to pre-train protein embeddings. Consistent with aforementioned results without pre-training, self-supervised pre-trainings of individual protein modalities differ in their strengths of predicting affinity or contacts. We further explore self-supervision on top of cross-modality learning, ask which pre-training scheme is beneficial in what circumstances of CPAC prediction, and provide conjectures to underlying reasons.

The rest of the manuscript is organized as follows. In Materials and Methods, we will start with our curated, labelled and unlabelled data, to supervise model training and pre-train protein embedding, respectively. After introducing a backbone model for CPAC prediction and our modifications, we will introduce our methods of cross-modality learning and multi-modal self-supervised learning. In Results, we will first examine performances from single- and multi-modal learning without pre-training. We will then examine self-supervised pretraining within and across modalities.

## 2 Materials and Methods

### 2.1 Data

#### Labelled Dataset

We evaluate compound–protein affinity and contact (CPAC) prediction methods through performing training and inference on a CPAC benchmark set (Karimi *et al.*, 2020; You and Shen, 2020) as follows.

##### (i) Data source

The diverse dataset contains 4,446 pairs between 1,287 proteins and 3,672 compounds that are collected from PDBbind (Liu *et al.*, 2015) and BindingDB (Liu *et al.*, 2007) together with their affinity labels. In addition, their contact labels are gathered from the corresponding cocrystal structures deposited in the PDBsum database (Laskowski *et al.*, 2018) using LigPlot. Histograms of protein and compound lengths, measured in the number of protein residues and that of compound atoms, are shown in Appendix A (Fig. S1).

##### (ii) Protein and compound graphs

No 3D structures of proteins or compounds are used. Instead, RaptorX-Contact (Xu, 2019) is used to predict contact maps of proteins from sequences, where evolutionary information from multiple sequence alignment and structural information from its labels are additionally included. Only binary contact maps are used without 3D structural information, thus called 2D graphs. RDKit (Landrum *et al.*, 2006) is used to convert 1D SMILES into 2D chemical structures for compounds, after sanitization.

##### (iii) Dataset split

The labelled dataset is split into subsets of various challenging levels in generalizability: 795 pairs involving unseen proteins (proteins not present in the training set), 521 pairs involving unseen compounds, and 205 for unseen both; whereas the rest is randomly split into training (2,334) including validation and the default test (591) sets (Karimi *et al.*, 2020). Statistics of the dataset split is presented in Table 1.

**Table 1.** Statistics of the dataset splits for affinity and contact prediction.

|  |  | 3,672 compounds |  |
| --- | --- | --- | --- |
|  |  | 3,100 | 572 |
| 1,287 proteins | 1,228 | Training set: 2,334 pairs | Unseen-compound test set: 521 pairs |
|  |  | Seen-both test set: 591 pairs |  |
|  | 59 | Unseen-protein test set: 795 pairs | Unseen-both test set: 205 pairs |

#### Unlabelled datasets

We pre-train protein embeddings using two unlabelled datasets of different scales. Both are from Pfam-A, a database of protein domain sequences (Mistry *et al.*, 2021): (i) The *smaller set* with ground-truth structure information consists of 60,137 sequences from Pfam-A with PDB entries (Berman *et al.*, 2000), from which we extract contact maps from their PDB structures (two residues are deemed in contact if their *C_β_*, or *C_α_* for glycines, are within 8Å). (2) The *larger set* not necessarily with ground-truth structure information is Pfam-A RP15 which consists of 12,798,671 sequences with 15% Representative Proteomes comembership (Chen *et al.*, 2011) threshold applied. Histograms of protein lengths are shown in Appendix A (Fig. S2).

### 2.2 Model Backbone

The backbone of a CPAC prediction model is a system that is given a compound–protein pair as inputs and simultaneously predicts intermolecular affinity and atom–residue contacts as outputs. Here we adopt the state-of-the-art CPAC model, DeepAffinity+ (Karimi *et al.*, 2020), as our models’ backbone.

Mathematically, given a compound–protein 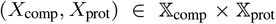 consisting of *N*_comp_ atoms in each compound and *N*_prot_ residues in each protein (padding is applied to ensure fixed sizes for all compounds or proteins), a CPAC model 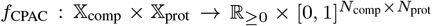 aims at predicting both the compound–protein affinity *z*_aff_ and the intermolecular atom–residue contacts *Z*_cont_. It includes the following three major components as shown in Figure 1.

1. **Neural-network encoders** 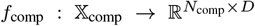 and 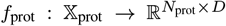 that separately extract embeddings ***H***_comp_ for the compound *X*_comp_ and ***H***_prot_ for the protein *X*_prot_ where *D* is the hidden dimension. In DeepAffinity+ the compounds are available in 2D chemical graphs and proteins are only available in 1D aminoacid sequences. Accordingly, DeepAffinity+ used graph neural networks (GNN) such as GCN and GIN (Kipf and Welling, 2016; Veličković *et al.*, 2017) to encode 2D chemical graphs of compounds and hierarchical recurrent neural network (HRNN) (El Hihi and Bengio, 1996) to encode 1D amino-acid sequences of proteins.
2. **Contact module** 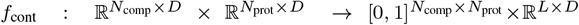 takes molecular embeddings from the encoders ***H***_comp_ and ***H***_prot_ as inputs, employs a joint attention mechanism (Karimi *et al.*, 2019, 2020) to output the atom–residue interaction matrix ***Z***_cont_, and jointly embeds the compound–protein pair into ***H***_cp_, where *L* is the hidden length determined by *N*_comp_ and *N*_prot_.
3. **Affinity module** *f*_aff_ ℝ^*L*×*D*^ → ℝ predicts the affinity *z*_aff_ given the joint embedding ***H***_cp_. It consists of 1D convolutional, pooling layers, and multi-layer perceptron (MLP). Note that the contact-predicting interaction module feeds the affinity module, making affinity prediction intrinsically interpretable by the underlying contacts.

**Fig.1.**
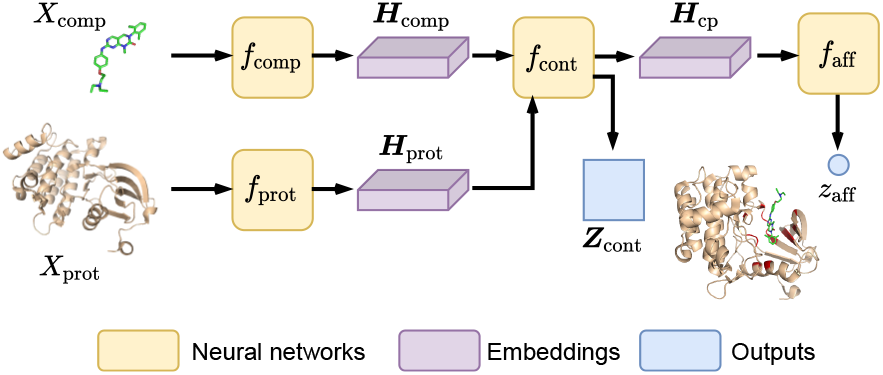
Illustration of thebackbone model fcPAC for compound–protein affinity and contact (CPAC) prediction.

After the CPAC model *f*_CPAC_ forwardly generates the outputs (*z*_aff_, ***Z***_cont_), true labels (*y*_aff_, ***Y***_cont_) are provided to calculate the loss, *l*_CPAC_, which consists of affinity loss *l*_aff_, intermolecular atom–residue contact loss *l*_cont_ and three structure-aware sparsity regularization losses *l*_group_, *l*_fused_, and *l*_L1_ as described in (Karimi *et al.*, 2020):

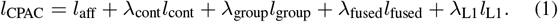

The model is trained end to end while the training loss is minimized. More details for the pipeline can be found in (Karimi *et al.*, 2020).

### 2.3 Single-Modality Protein Embeddings

In the conventional structure-free CPAC pipeline, compounds are represented as 2D chemical graphs since 1D SMILES strings have limited descriptive power and known worse performance in many tasks (Karimi *et al.*, 2020, 2019; Li *et al.*, 2020), whereas proteins are usually represented as 1D amino-acid sequences without exploration of other modalities. We delve into this under-explored area, proposing to utilize multi-modality protein data for CPAC prediction.

#### 1D sequences

We follow DeepAffinity+ (Karimi *et al.*, 2020) as described in Section 2.2 and use HRNN to encode protein sequences. One change we made is replacing the hierarchical joint attention with naïve joint attention in the interaction module expressed as:

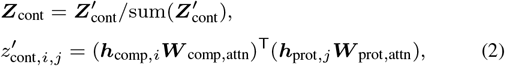

where *z_i,j_* = ***Z*** [*i,j*], ***h**_i_* = ***H***[*i*, :], *i* = 1,…, *N*_comp_, *j* = 1,…,*N*_prot_; ***W***_comp,attn_ and ***W***_prot,attn_ are two learnable attention matrices.

#### 2D contact maps

We propose to adopt the 2D modality of proteins as additional inputs and model them as graphs with the following reasons. (1) Graphs are more structure-aware compared to 1D sequences, potentially resulting in better generalizability. (2) Graphs are concise yet informative (focusing on pairwise residue interactions) compared to the data structure of 3D coordinates (which are also harder to predict than contact maps) (Cao and Shen, 2020). (3) The recent surge of models for graph learning (Kipf and Welling, 2016; Veličković *et al.*, 2017) provides advanced tools to facilitate graph representation learning.

As unbound or ligand-bound structure data is not readily available for many proteins, we use sequence-predicted 2D contact maps (Xu, 2019) and can also use AlphaFold2 (Jumper *et al.*, 2021). Thereby, we additionally represent a protein input *X*_prot_ as a graph 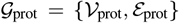 where vertices stand for residues and edges exist between residues predicted to be in contact. The graphs are associated with feature matrix ***F***_prot_ ∈ ℝ^*N*_prot_×*D*^ (embedded amino-acid types of residues) and the adjacency matrix ***A***_prot_ ∈ {0, 1}^*N*_prot_×*N*_prot_^ (binary contact map). We employ an expressive GNN model, graph attention network (GAT) (Veličković *et al.*, 2017) with *K* layers as the protein encoder *f*_prot_ to extract graph embeddings, with the formulation of each layer’s forward propagation as:

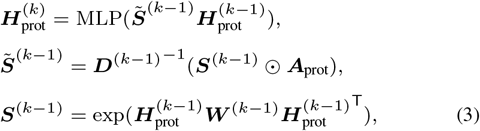

where 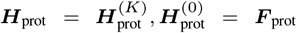, ⊙ is the element-wise multiplication, ***J***_*N*_prot,1__ is an all-ones matrix with size *N*_prot_ × 1, and ***W***^(*k*–1)^ isa learnable weight matrix. Comparison with the simplest GNN model, graph convolutional network (GCN) is conducted in Appendix B to demonstrate the necessity of adopting the more expressive GAT.

### 2.4 Cross-Modality Protein Embeddings

To integrate the knowledge from both 1D and 2D protein modalities, we introduce two cross-modality protein embedding schemes as follows.

#### Cross-modality concatenation

A simple integration model is to concatenate the extracted embeddings of the 1D and 2D modalities encoded by HRNN and GAT, respectively, as shown in Figure 2(a). Indeed, concatenation is commonly used in previous work (Hamilton *et al.*, 2017; Xu *et al.*, 2018) to preserve information from different sources. The concatenated output is fed to a multi-layer perception (MLP) for the final protein embedding ***H***_prot_.

**Fig. 2.**
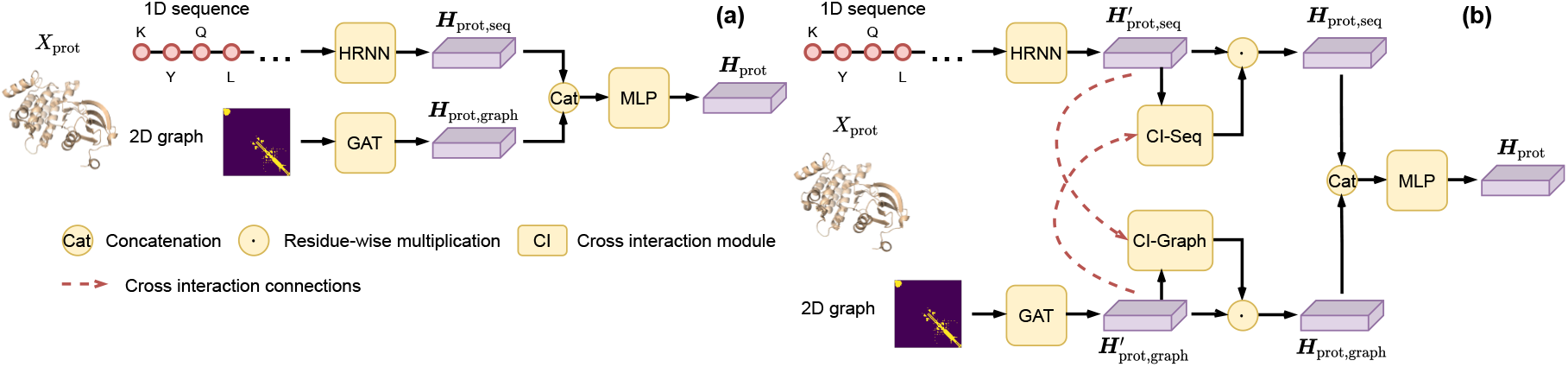
Cross-modality encoders for proteins (*f*_prot_ in Figure 1) to capture and integrate knowledge across data modalities. (a) Naïve concatenation preserves information from different sources, and (b) cross interaction additionally introduces information flows between modalities.

#### Cross-modality cross interaction

Although the aforementioned concatenation strategy preserves the information of individual modalities, the encoding processes for the two modalities are isolated. In other words, the two types of embeddings from different modalities were independently encoded and then mixed through concatenation. However, the different modalities of proteins are intrinsically correlated with each other and could be coupled in a properly-designed representation-learning process. Therefore, we introduce a cross interaction module to facilitate the encoder to learn protein embeddings from correlated data (1D and 2D modalities), as shown in Figure 2(b). Specifically, given the outputs of encoders 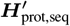 and 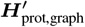, we calculate sequence and graph cross-modality outputs ***H***_prot,seq_ and ***H***_prot,graph_, respectively:

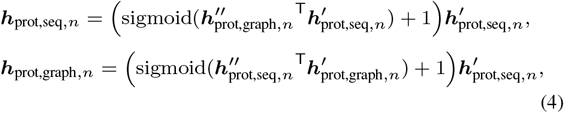

where 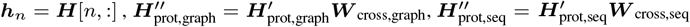; ***W***_cross,seq_ and ***W***_cross,graph_ are learnable weights.

Instead of independently extracting knowledge from protein modalities (1D sequences and 2D contact maps), the cross interaction module enforces a learned relationship between the encoded embeddings of the two protein modalities, which is expected to better capture the information from the correlated modalities and to benefit the affinity and contact prediction. Again, ***H***_prot,seq_ and ***H***_prot,graph_ (now with information from each other) are concatenated and fed to an MLP for the final protein embedding ***H***_prot_.

The idea of cross interaction was previously introduced in (Tan and Bansal, 2019) and modified here as follows. (1) We do not normalize cross interaction along residues (sequence length is 1,000 here) since it would significantly change the scale of the residue embeddings. (2) We restrict the cross interaction for each residue in the range of [0, 1] with sigmoid function to represent the cross-modality “interaction strength”.

### 2.5 Multi-Modality Self-Supervised Pre-Training

On top of the aforementioned cross-modality learning models, we further propose self-supervised pre-training for the following two reasons. (1) The paired and labelled data curated for CPAC (Karimi *et al.*, 2020) are limited (4,446 compound–protein pairs in total), while there are more than billions of unpaired and unlabelled data available (here we make use of protein domain sequences as described in Section 2.1). Exploiting such abundant unlabelled data would generate context-relevant embeddings for downstream, as previously explored under unsupervised learning in CPAC prediction (Karimi *et al.*, 2019). (2) Compared to conventional unsupervised learning, recently emerging self-supervised learning on both sequences (Devlin *et al.*, 2018) and graphs (You *et al.*, 2020b,a, 2021, 2022) further exploits the benefit from unlabelled data.

For reasons above we introduce the following pretraining strategies, as illustrated in Figure 3. In addition, graph contrastive learning GraphCL (You *et al.*, 2021) is also applied.

**Fig. 3.**
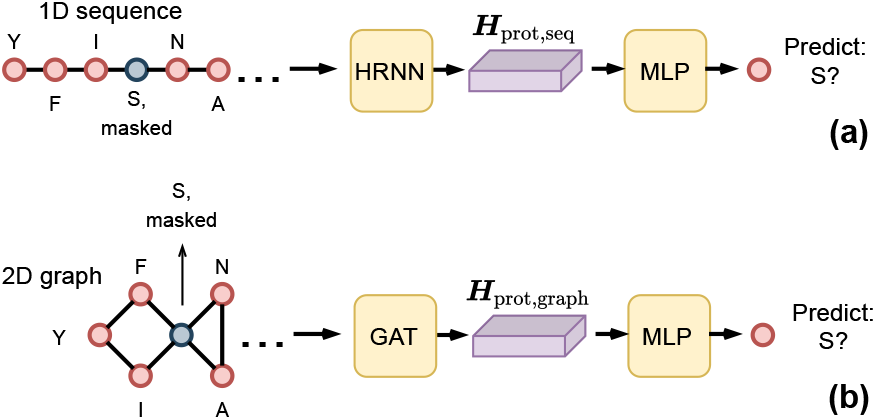
Self-supervised tasks for pre-training cross-modality encoders (Figure 2) in CPAC. (a) Masked language modeling (MLM) takes the randomly masked amino-acid sequences as inputs, predicting the masked residues with network outputs, and (b) graph completion (GraphComp) with inputting masked-residues contact maps, makes prediction for the masked tokens.

#### Masked language modeling for sequences

We adopt masked language modeling (MLM) for the 1D sequence encoder HRNN, which is well-known as the dominant pre-training strategy in natural language processing (Devlin *et al.*, 2018). MLM takes the randomly masked aminoacid sequences as inputs, and tries to predict the masked residues (we use residue types for a proper self-supervising “curriculum”) with network outputs, as illustrated in Figure 3(a). The mathematical formulation of MLM optimization is expressed as:

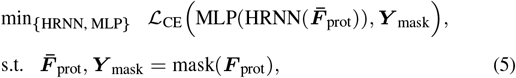

where 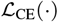 is cross-entropy loss, 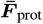 is the masked feature matrix, ***Y***_mask_ is the masked residues, and mask(·) is the masking function.

MLM reconstructs and enforces the missing knowledge through utilizing the sequential relation (where the information flow is specified by sequential inputs), which aligns with the 1D-modality model exploiting protein sequence information. We thus hypothesize that MLM pre-training provides performance gains in the tasks where the 1D-modality model has performed well, i.e. affinity prediction, which is supported by experimental results in Section 3.4.

#### Masked graph modeling (graph completion) for contact maps

Self-supervision on graph-structured data recently raises great interests with numerous self-supervised tasks proposed (You *et al.*, 2020b,a, 2021). We choose a simple and effective scheme, graph completion or GraphComp (You *et al.*, 2020b), to pre-train the 2D graph encoder GAT. GraphComp can be viewed as “the graph version of MLM”: it takes graphs with randomly masked residues as input and aims at making prediction for the masked tokens using the structure-aware graph information, as illustrated in Figure 3(b). GraphComp optimization is mathematically formulated as:

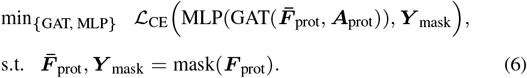

#### Joint self-supervised pre-training

Besides single-modality pretraining, we also propose joint pre-training for the cross-modality models, that simultaneously performs MLM and GraphComp for self-supervision (since sequence and protein encoders share the amino-acid embedding layer, we cannot individually pre-train them and then load the checkpoints). Given benefits from single-modality pre-training, we expect more benefits can be achieved from multi-modality pre-training in both tasks of affinity prediction (where 1D modality models performed well) and contact prediction (where 2D modality models performed well). Results in Section 3.5 partly justified the added benefits.

Details about model training, including hyperparameters, are in Appendix H.

## 3 Results and Discussion

We organize results and discussion as follows. Experiments on cross-modality protein embeddings are presented in Sections 3.1 and 3.2, with additional generalizability tests and case studies. Self-supervised pretraining experiments on top of cross-modality models are reported in Sections 3.4 and 3.5.

### 3.1 Individual modalities have strengths in different tasks

Without pre-training, Table 2 reports various models’ performances for affinity prediction and contact predictionfor various test sets. Figure 4 further splits unseen molecules into proteins and compounds of different similarity bins compared to the training set.

**Fig. 4.**
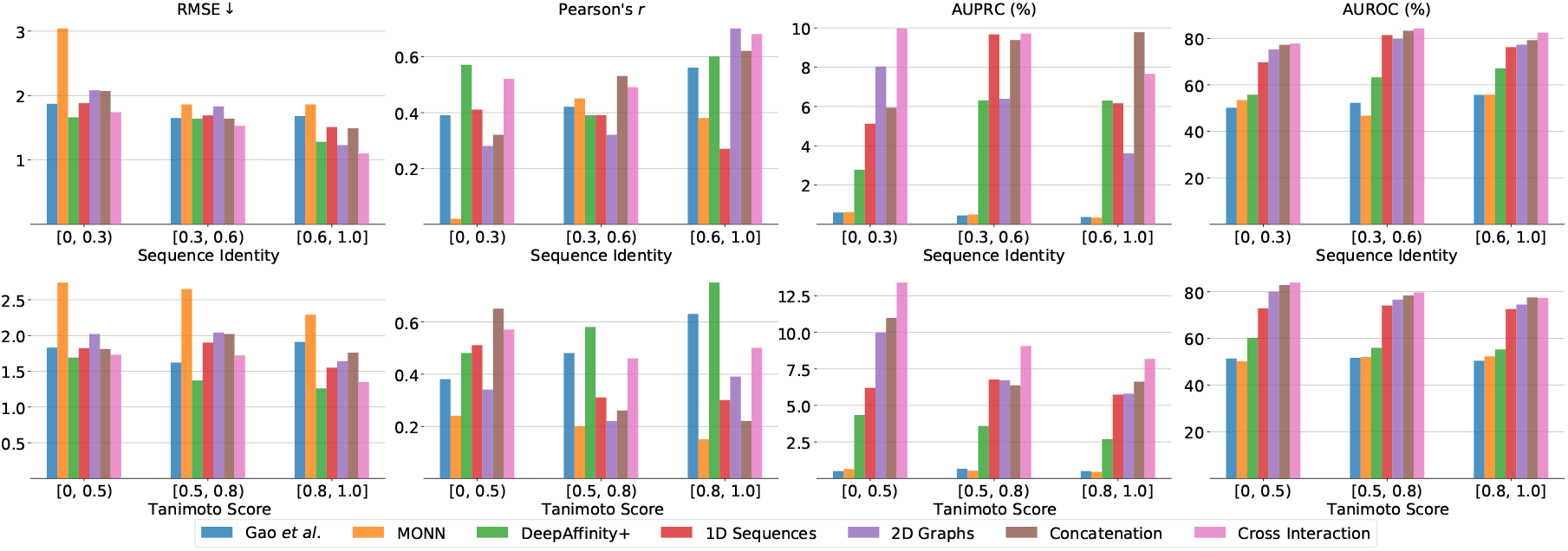
Generalizability test on various methods for predicting affinity (measured in RMSE and r) and contact (measured in AUPRC and AUROC).

**Table 2.**
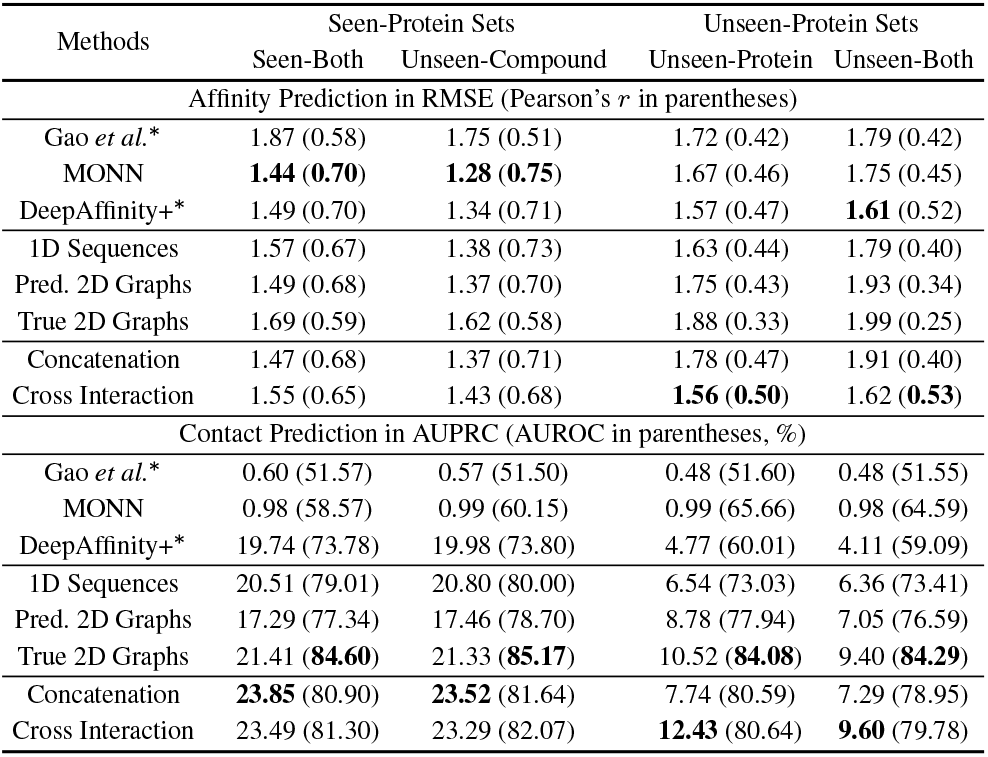
Comparison among competing methods and ours in compound–protein affinity prediction (measured by RMSE and Pearson’s correlation coefficient *r*) and contact prediction (measured by AUPRC and AUROC). * denotes the cited performances. Boldfaced numbers are the best performances for given test sets. We note that, as intermolecular contacts only represent a minority (around 0.4%) of all compound–protein atom-residue pairs, AUPRC is a much more relevant measure than AUROC for assessing contact prediction.

In affinity prediction, 1D sequences or 2D graphs did not lead to significant difference for seen proteins. However, speaking of unseen proteins or even non-homologous proteins (sequence identity below 30%) where model generalizability is required, 1D sequences dominated over 2D graphs as inputs for affinity prediction (0.1 lower in RMSE).

One conjecture is that the information in graphs might be more difficult to learn compared to sequences (the training RMSE losses are 0.71 and 0.99 for 1D and 2D modalities, respectively). Moreover, affinity prediction for unseen-protein cases are not as challenging as intermolecular contact prediction to show the benefit of the 2D modality (shown next), as contact prediction often involves tens of thousands of values (rather than a single value) to fit for each compound–protein pair.

In contact prediction, encoding proteins as 1D sequences again performed better (+3.22% at AUPRC and +1.67% at AUROC) for seen proteins (the proteins in the training set). However, encoding 2D contact maps (graphs) significantly outperformed doing 1D protein sequences (+4.91% at AUPRC and +2.24% at AUROC) for unseen proteins (Table 2) and even more for non-homologous proteins (Figure 4). Using “true” contact maps from (unbound) protein structures showed the same and improved AUROC.

We conjecture that sequential knowledge encoded in 1D amino-acid sequences is well captured especially for seen proteins after training. The sequential dependency learned from the encoder could be accurate toward intermolecular contact prediction for close or even distant homologs of seen proteins. However such dependency is less generalizable to unseen or non-homologous proteins. In contrast, the structural topology information encoded in protein 2D contact maps is more difficult for graph neural networks to capture even for seen proteins, leading to the worse contact predictions for seen proteins. But the information can generalize to unseen proteins well toward contact prediction. In particular, even when sequence similarity for non-homologous proteins (to training ones) is too low to be detectable using RNNs, binding-pocket (subgraph) similarity could still preserve and be detected in 2D contact maps using GNNs thus eventually leads to much better intermolecular contact prediction (Figure 4).

### 3.2 Cross-modality models combine the strengths

Fusing two modalities’ knowledge together, even by a simple concatenation strategy, could get the best of both modalities. Specifically, the cross-modality model by concatenation had better contact prediction than single-modality models (Table 2). It also had a boost in affinity prediction (better than the 2D single-modality model and slightly worse than the 1D single-modality model).

Enforcing a learned correlation between the 1D and 2D embeddings rather than independently learning two individual embeddings, the cross-modality model with cross interaction further improved affinity prediction and actually had the best affinity accuracy among all methods for unseen proteins or unseen both. Moreover, it impressively achieved the best AUPRC for unseen proteins and unseen both. These results re-enforce our rationale that the learned correlation between embeddings from different modalities can better capture the data and better perform CPAC predictions.

Our models compare favorably to the state-of-the-art (SOTA) models. They used similar backbone as DeepAffinity+ (Karimi *et al.*, 2020) and revised the joint attention mechanism as mentioned in Section 2.4; thus our 1D sequence-based single-modality model and DeepAffinity+, both using protein sequences, had similar performances in affinity prediction but ours improved contact prediction. Our cross-modality models further improved the performance compared with SOTAs including Gao *et al.* (after being converted from a binary predictor) (Gao *et al.*, 2018), MONN (Li *et al.*, 2020) and DeepAffinity+ (Karimi *et al.*, 2020), especially for unseen proteins (Table 2) and non-homologous proteins (Figure 4).

When the protein sequence encoder was changed from HRNN to a pre-trained Transformer, no improvement was found (Appendix C).

### 3.3 Case studies for cross-modality models

All methods are compared in five case studies about compound–protein pairs (Karimi *et al.*, 2020). With detailed results included in Appendix D, we conclude that one or both cross-modality models improved over DeepAffinity+ in AUPRC for four of the five cases. They performed on par with DeepAffinity+ in the precision of the predicted top-10 contacts. The case of LHL–LCK presented the most improvement in the precision of top-10 predicted contacts, from 0.4 to 0.6, as visualized in Figure 5.

**Fig. 5.**
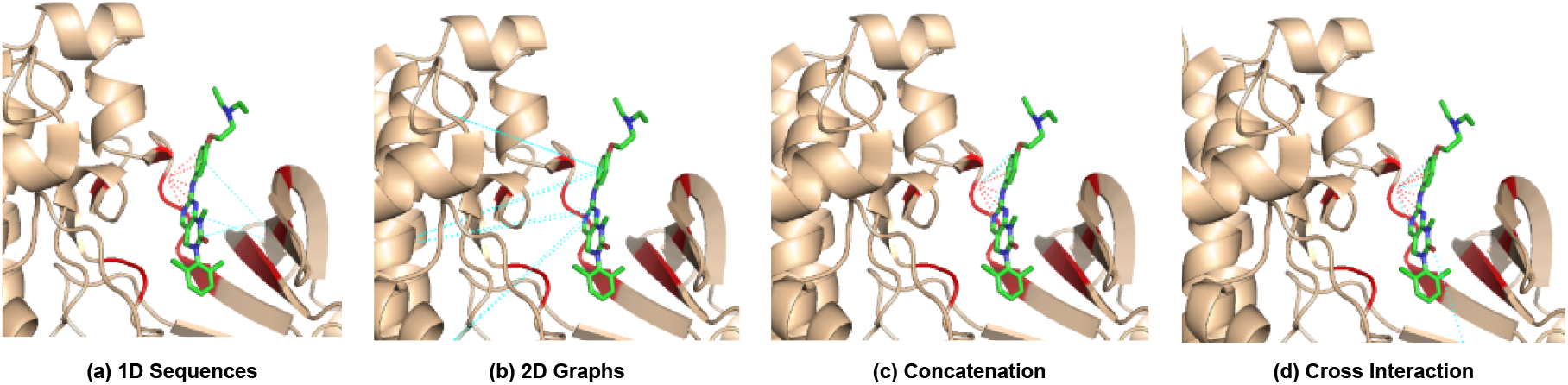
Visualizing top-10 atom–residue contacts predicted by single- and cross-modal learning for the compound–protein pair of LHL–LCK. Compounds are shown in sticks (green for carbon, red for oxygen and blue for nitrogen atoms), proteins in wheat cartoons (with red patches of binding sites), and predicted contacts in dashed lines (red for true positives and cyan for false positives).

### 3.4 Single-modality pretraining further enhance individual modalities’ strengths

We proceed to pre-train our cross-modality model (cross interaction) in a single-modality setting. In other words, we pretrain the protein sequence and graph encoders using MLM and GraphComp, respectively. The results are detailed in Table 3.

**Table 3.** Comparison among different pre-training settings (masked language modeling and graph completion, with graph contrastive learning in Appendix E) based upon the cross interaction model in compound–protein affinity and contact prediction. Boldfaced are the best performances.

| Cross Interaction | Seen-Protein Sets |  | Unseen-Protein Sets |  |
| --- | --- | --- | --- | --- |
|  | Seen-Both | Unseen-Compound | Unseen-Protein | Unseen-Both |
| Affinity Prediction in RMSE (Pearson's $r$ in parentheses) | | | | |
| Non Pre-Train | 1.57 (0.66) | 1.46 (0.68) | 1.63 (0.49) | 1.64 (0.54) |
| MLM-S | **1.53 (0.64)** | **1.40 (0.68)** | **1.46 (0.56)** | **1.53 (0.58)** |
| GraphComp-S | 1.62 (0.59) | 1.44 (0.66) | 1.60 (0.43) | 1.67 (0.47) |
| MLM+GraphComp-S | 1.64 (0.58) | 1.46 (0.65) | 1.65 (0.39) | 1.65 (0.50) |
| MLM-L | 1.59 (0.62) | 1.46 (0.65) | 1.62 (0.47) | 1.63 (0.57) |
| MLM+GraphComp-L | 1.58 (0.62) | 1.45 (0.66) | 1.74 (0.33) | 1.85 (0.32) |
| Contact Prediction in AUPRC (AUROC in parentheses, %) |  |  |  |  |
| Non Pre-Train | 23.91 (79.48) | 23.06 (80.60) | 11.40 (77.73) | 8.41 (76.42) |
| MLM-S | 23.78 (80.34) | 23.33 (81.09) | 7.73 (77.44) | 6.44 (76.42) |
| GraphComp-S | 23.63 (79.71) | 23.41 (81.31) | 11.36 (76.67) | 9.36 (76.00) |
| MLM+GraphComp-S | **24.13 (82.09)** | **23.65 (82.70)** | 11.38 (78.75) | 10.83 (78.63) |
| MLM-L | 23.30 (80.40) | 23.05 (81.18) | 11.35 (81.01) | 9.40 (79.46) |
| MLM+GraphComp-L | 23.71 (81.21) | 23.22 (82.33) | **13.47 (82.00)** | **11.17 (80.10)** |

Different pre-training strategies showed different performances relative to no pre-training, depending on the task (affinity or contact prediction) and the test set (seen or unseen proteins/compounds). Consistent with our earlier observation of single-modality models without pretraining, pre-training the embedding of a single modality tended to enhance the strength of the corresponding modality. Specifically, sequence pretraining with MLM, especially with the smaller unlabelled protein dataset, improved upon what the 1D protein modality is good at — affinity prediction, for unseen proteins. MLM over the larger unlabelled set of protein sequences did not show much more benefits, possibly due to the fact that the smaller unlabelled set and the labelled test sets are biased with protein of structures. Meanwhile, graph pretraining with GraphComp, over the smaller or the larger unlabelled protein dataset, improved upon what the 2D protein modality is good at – contact prediction, mainly for unseen both. Replacing GraphComp (You *et al.*, 2020b) with contrastive learning (GraphCL) (You *et al.*, 2021) had similar performances (Appendix E).

We observe some trade-off between affinity and contact prediction while pre-training a single modality. Part of the reason could be that the two tasks compete with each other while their weighted losses are summed together. The question that remains is whether and how the pre-training strategies for individual modalities can be combined to further enhance model accuracy and generalizability, which is addressed next.

### 3.5 Multi-modal joint pre-training could further synergize 1D and 2D modalities

We further pretrain our cross-modality model in a multi-modal setting. In other words, we jointly pretrain both the sequence and the graph encoders that share layers. The results are reported in Table 3 as before.

We found that jointly pre-training sequence and graph embedding with the smaller unlabelled dataset didn’t change affinity prediction much for unseen proteins and improved contact prediction for the most challenging case of unseen both (+2.4% in AUPRC compared to no pretraining). Interestingly, doing so with the larger unlabelled dataset again improved contact prediction for the most challenging case of unseen both (+2.7% in AUPRC compared to no pretraining) and additionally did so for the unseen proteins (+2.1% in AUPRC compared to no pretraining). Impressively, the joint pre-training strategies with predicted protein contact maps even outperformed non pre-training with actual protein contact maps. In the end, the cross-modality model (cross interaction) with joint sequence-graph pretraining over the larger set achieved the best contact prediction for both unseen proteins and unseen both. And doing that over the smaller set achieved best balanced improvement in affinity and contact prediction, potentially suggesting the importance of data quality over data quantity.

We also tested additional pre-training for embedding 2D compound graphs on top of the cross-modality model with joint pretraining of protein data. To do so, we leveraged unlabelled compound data from STITCH. Further improvements, albeit moderate, were observed (Appendix G).

## 4 Conclusion

In this paper, we address two major challenges to advance explainable prediction of compound–protein affinity (or CPAC, compound–protein affinity and contact): the sequence-dominant yet structure-naive models and the scarce labelled data. By introducing multi-modal and selfsupervised learning for the first time to CPAC prediction, we address both challenges through fostering context- and task-relevant protein embedding. Specifically, to overcome structure naivety, we treat protein data as available in both modalities of 1D sequences and 2D graphs (predicted) and introduce cross-modality learning for sequence- and structure-aware protein embeddings. Empirical results indicated that individual modalites excel in different tasks and our approach of cross-modality learning could bring out the best of both modalities. Additionally, to overcome labelled-data scarcity, we design self-supervised learning strategies within and across modalities to pretrain cross-modal protein embedding. Empirical results indicated that cross-modal learning with joint pre-training can further improve model generalizability for unseen molecules and outperform the state of the art. Meanwhile, there is still much to do for improving the synergy between both tasks of affinity and contact prediction.

## Acknowledgements

Part of the computing resources are provided by Texas A&M University High Performance Research Computing (HPRC).

