## supplement for "Cross-Modality and Self-Supervised Protein Embedding for Compound–Protein Affinity and Contact Prediction"

### A Dataset Statistics

Histograms of protein and compound lengths of the evaluation dataset are shown in Figure S1, and histograms of protein lengths of the pre-training datasets are shown in Figure S2.

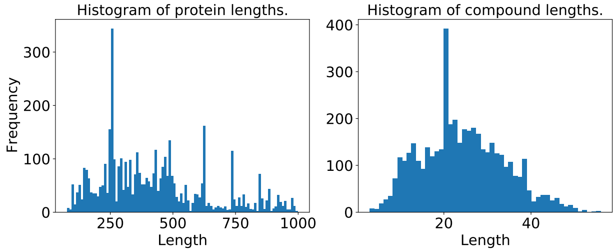


**Fig. S1.** Histograms of protein and compound lengths in the compound-protein affinities and contacts prediction dataset.

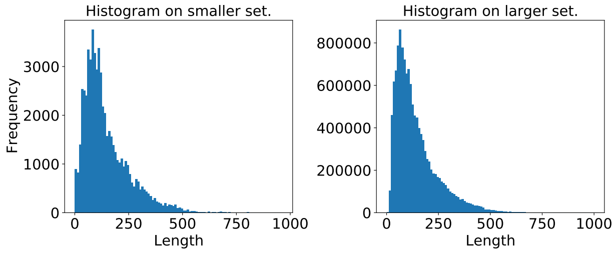


**Fig. S2.** Histograms of protein lengths in the pre-training datasets.

### B Comparison between GAT and GCN for protein-graph embedding

Experiments on comparing GAT and GCN on single-modality 2D graph model and two cross-modality models are presented in Tables S1 and S2.

### C Comparison between HRNN and Transformer for protein-sequence embeding

Experiments on comparing HRNN and Transformer on single-modality 1D sequence model and two cross-modality models are presented in Table S3.

### D Case Studies

Five case studies on compound-protein pairs including AL1-CA2, IT2-CA2, CPB-PYGM, T68-PYGM and LHL-LCK are shown in Table S4 and visualizations in Figures S3-S7. For detailed case description please refer to (Karimi *et al.*, 2020).

Table S1. Comparison between GAT and GCN in compound-protein affinities prediction (measured by RMSE and Pearson's correlation coefficient).

| Methods | | GAT | | GCN | | | GAT | | GCN | |
|---|---|---|---|---|---|---|---|---|---|---|
| | | S.-Both | U.S.-Comp. | S.-Both | U.S.-Comp. | | U.S.-Prot. | U.S.-Both | U.S.-Prot. | U.S.-Both |
| Single Modality | RMSE | 1.49 | 1.37 | 1.59 | 1.38 | RMSE | 1.75 | 1.93 | 1.66 | 1.75 |
| (2D Graphs) | Pearson's $r$ | 0.68 | 0.70 | 0.61 | 0.68 | Pearson's $r$ | 0.43 | 0.34 | 0.41 | 0.41 |
| Cross Modality | RMSE | 1.47 | 1.37 | 1.51 | 1.40 | RMSE | 1.78 | 1.91 | 1.56 | 1.61 |
| (Concatenation) | Pearson's $r$ | 0.68 | 0.71 | 0.66 | 0.68 | Pearson's $r$ | 0.47 | 0.40 | 0.46 | 0.51 |
| Cross Modality | RMSE | 1.55 | 1.43 | 1.51 | 1.33 | RMSE | 1.56 | 1.62 | 1.61 | 1.70 |
| (Cross Interaction) | Pearson's $r$ | 0.65 | 0.68 | 0.66 | 0.71 | Pearson's $r$ | 0.50 | 0.53 | 0.46 | 0.47 |

Table S2. Comparison between GAT and GCN in compound-protein contacts prediction (measured by AUPRC and AUROC).

| Methods | | GAT | | GCN | | | GAT | | GCN | |
|---|---|---|---|---|---|---|---|---|---|---|
| | | S.-Both | U.S.-Comp. | S.-Both | U.S.-Comp. | | U.S.-Prot. | U.S.-Both | U.S.-Prot. | U.S.-Both |
| Single Modality | AUPRC (%) | 17.29 | 17.46 | 11.14 | 10.70 | AUPRC (%) | 8.78 | 7.05 | 11.65 | 9.50 |
| (2D Graphs) | AUROC (%) | 77.34 | 78.70 | 74.19 | 74.15 | AUROC (%) | 77.94 | 76.59 | 79.36 | 78.45 |
| Cross Modality | AUPRC (%) | 23.85 | 23.52 | 21.88 | 22.06 | AUPRC (%) | 7.74 | 7.29 | 7.51 | 6.62 |
| (Concatenation) | AUROC (%) | 80.90 | 81.64 | 80.74 | 81.15 | AUROC (%) | 80.59 | 78.95 | 77.28 | 77.58 |
| Cross Modality | AUPRC (%) | 23.49 | 23.29 | 23.40 | 23.51 | AUPRC (%) | 12.43 | 9.60 | 7.11 | 6.77 |
| (Cross Interaction) | AUROC (%) | 81.30 | 82.07 | 81.39 | 82.03 | AUROC (%) | 80.64 | 79.78 | 74.93 | 76.29 |

Table S3. HRNN vs Transformer.

| Tasks | | HRNN | | Transformer | | | HRNN | | Transformer | |
|---|---|---|---|---|---|---|---|---|---|---|
| | | S.-Both | U.S.-Comp. | S.-Both | U.S.-Comp. | | U.S.-Prot. | U.S.-Both | U.S.-Prot. | U.S.-Both |
| Affinity | RMSE | 1.57 | 1.38 | 1.69 | 1.49 | RMSE | 1.63 | 1.79 | 1.88 | 1.98 |
| prediction | Pearson's $r$ | 0.67 | 0.73 | 0.59 | 0.66 | Pearson's $r$ | 0.44 | 0.40 | 0.34 | 0.36 |
| Contact | AUPRC (%) | 20.51 | 20.80 | 12.09 | 11.76 | AUPRC | 6.54 | 6.36 | 0.62 | 0.58 |
| prediction | AUROC (%) | 79.01 | 80.00 | 64.25 | 63.34 | AUROC | 73.03 | 73.41 | 51.99 | 52.01 |

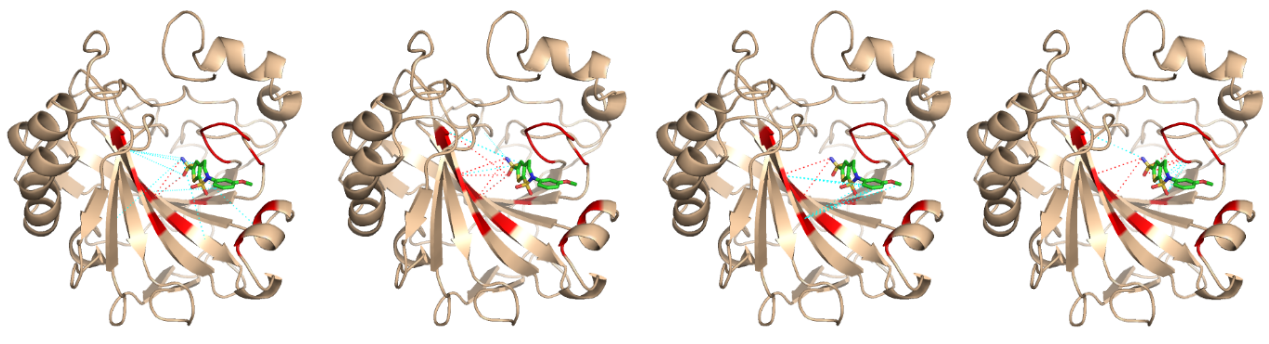

(a) 1D Sequences (b) 2D Graphs (c) Concatenation (d) Cross Interaction

**Fig. S3.** Compound-protein pair AL1-CA2 top-10 contacts prediction visualization. The dashed lines in red and pale cyan highlight correct and incorrect predictions, respectively, according to native, direct contacts retrieved by LigPlot.

### E Comparing GraphComp and GraphCL for pretraining protein graphs

Experiments on comparing GraphComp (You *et al.*, 2020b) and GraphCL (You *et al.*, 2020a) on single- modality pretraining for embedding 2D graphs are reported in Table S5.

### F Pre-Training on the Larger Set of Unlabelled Proteins at Different Epochs

Figures S8 - S15 show the results of cross-modality models with sequence pre-training on the larger set for different epochs.

Table S4. Affinities and contacts prediction on five compound-protein pairs, AL1–CA2, IT2–CA2, CPB–PYGM, T68–PYGM and LHL–LCK.

| Methods | Affinity Error↓ | AUPRC (%) | AUROC (%) | Top-10 Contact Precision |
|---|---|---|---|---|
| | | AL1–CA2 | | |
| Gao et al. | 3.28 | 0.6 | 50.0 | 0.0 |
| DeepAffinity+ | 1.89 | 28.4 | 65.8 | 0.5 |
| 1D Sequences | 2.81 | 8.60 | 75.41 | 0.2 |
| 2D Graphs | 3.86 | 14.56 | 82.11 | 0.5 |
| Concatenation | 3.64 | 7.94 | 90.99 | 0.2 |
| Cross Interaction | 3.10 | 25.44 | 81.95 | 0.5 |
| | | IT2–CA2 | | |
| Gao et al. | 3.09 | 0.9 | 63.0 | 0.0 |
| DeepAffinity+ | 2.92 | 3.4 | 60.1 | 0.3 |
| 1D Sequences | 3.37 | 4.37 | 75.96 | 0.1 |
| 2D Graphs | 4.82 | 1.02 | 77.78 | 0.0 |
| Concatenation | 6.27 | 8.73 | 96.82 | 0.1 |
| Cross Interaction | 5.22 | 2.25 | 82.19 | 0.0 |
| | | CPB–PYGM | | |
| Gao et al. | 0.61 | 0.1 | 52.2 | 0.0 |
| DeepAffinity+ | 0.10 | 0.6 | 55.2 | 0.1 |
| 1D Sequences | 0.06 | 3.01 | 76.25 | 0.0 |
| 2D Graphs | 0.73 | 14.85 | 74.19 | 0.4 |
| Concatenation | 0.46 | 5.60 | 79.22 | 0.0 |
| Cross Interaction | 0.23 | 4.16 | 83.81 | 0.1 |
| | | T68–PYGM | | |
| Gao et al. | 1.80 | 0.6 | 63.5 | 0.0 |
| DeepAffinity+ | 0.68 | 67.5 | 94.4 | 1.0 |
| 1D Sequences | 0.99 | 63.08 | 91.78 | 1.0 |
| 2D Graphs | 0.24 | 52.20 | 90.82 | 1.0 |
| Concatenation | 0.51 | 77.73 | 96.73 | 1.0 |
| Cross Interaction | 0.94 | 73.10 | 97.62 | 1.0 |
| | | LHL–LCK | | |
| Gao et al. | 2.89 | 0.5 | 54.0 | 0.0 |
| DeepAffinity+ | 2.12 | 5.3 | 50.0 | 0.4 |
| 1D Sequences | 0.83 | 20.40 | 78.53 | 0.5 |
| 2D Graphs | 2.02 | 0.86 | 0.77 | 0.0 |
| Concatenation | 1.60 | 21.27 | 85.80 | 0.6 |
| Cross Interaction | 1.51 | 20.59 | 83.72 | 0.6 |

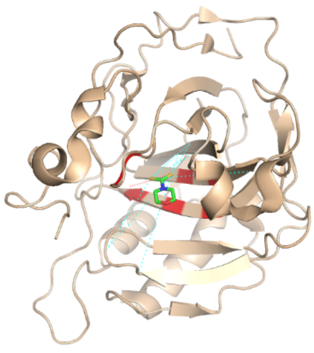

(a) 1D Sequences

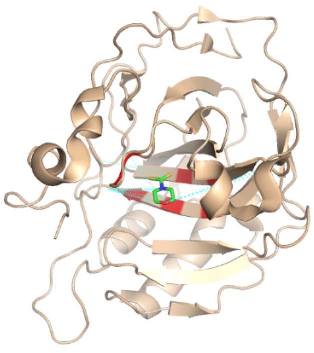

(b) 2D Graphs

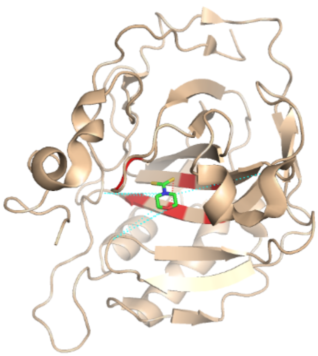

(c) Concatenation

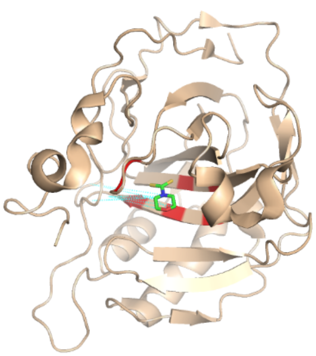

(d) Cross Interaction

**Fig. S4.** Compound-protein pair IT2-CA2 top-10 contacts prediction visualization.

### G Pre-Training for Compound Graph Embeddings

Experimental results on pre-training the compound graph encoder (GraphComp is performed here) on the cross interaction model with MLM+GraphComp pre-training for protein embeddings are shown in

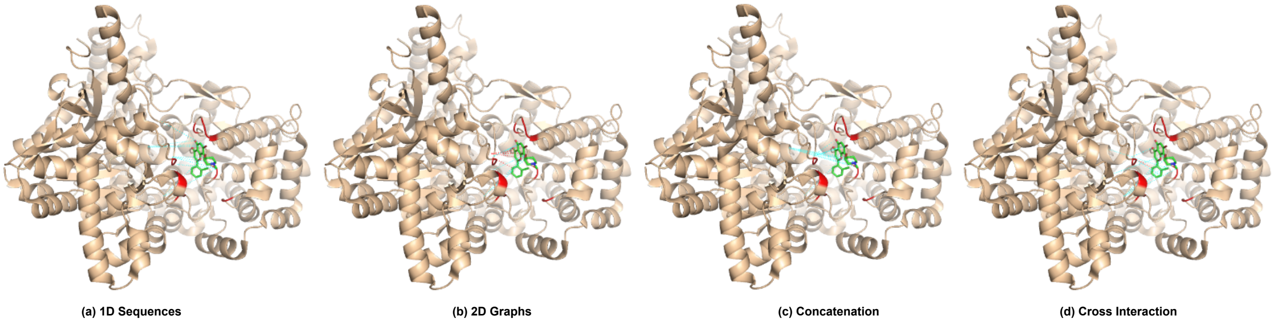


**Fig. S5.** Compound-protein pair CPB-PYGM top-10 contacts prediction visualization.

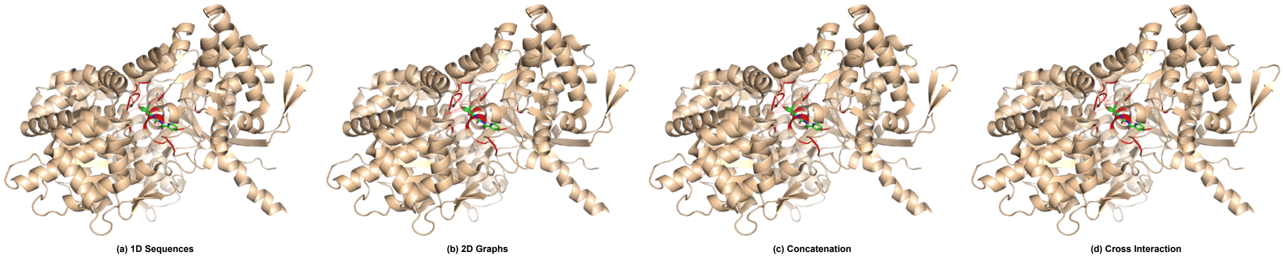


**Fig. S6.** Compound-protein pair T68-PYGM top-10 contacts prediction visualization.

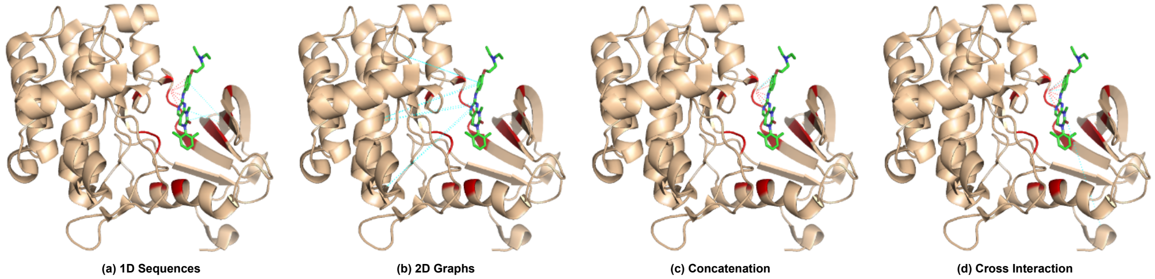


**Fig. S7.** Compound-protein pair LHL-LCK top-10 contacts prediction visualization.

Table S5. GraphComp vs GraphCL.

| Tasks | | GraphComp | | GraphCL | | | GraphComp | | GraphCL | |
|---|---|---|---|---|---|---|---|---|---|---|
| | | S.-Both | U.S.-Comp. | S.-Both | U.S.-Comp. | | U.S.-Prot. | U.S.-Both | U.S.-Prot. | U.S.-Both |
| Affinity prediction | RMSE | 1.62 | 1.44 | 1.53 | 1.41 | RMSE | 1.60 | 1.67 | 1.66 | 1.77 |
| | Pearson's $r$ | 0.59 | 0.66 | 0.67 | 0.70 | Pearson's $r$ | 0.43 | 0.47 | 0.44 | 0.46 |
| Contact prediction | AUPRC (%) | 23.63 | 23.41 | 18.15 | 17.30 | AUPRC | 11.36 | 9.36 | 11.09 | 8.55 |
| | AUROC (%) | 79.71 | 81.31 | 76.04 | 75.96 | AUROC | 76.67 | 76.00 | 73.46 | 70.94 |

Table S6. The pre-training compounds are collected from STITCH database (Szklarczyk *et al.*, 2016) with the same preprocess procedure conducted in (Karimi *et al.*, 2020).

### H Modeling Training

When no pre-training is performed, supervised models are trained over the training set of the labelled data and makes inference on the four test sets described in Section 2.1. We train models end-to-end with the following

Table S6. Comparison between w/ and w/o compound graph embedding pre-training (GraphComp) on the cross interaction model with MLM+GraphComp pre-training for protein embeddings.

| Methods | | w/o Comp. P.T. | | w/ Comp. P.T. | | | w/o Comp. P.T. | | w/ Comp. P.T. | |
|---|---|---|---|---|---|---|---|---|---|---|
| | | S.-Both | U.S.-Comp. | S.-Both | U.S.-Comp. | | U.S.-Prot. | U.S.-Both | U.S.-Prot. | U.S.-Both |
| Affinities | RMSE | 1.68 | 1.45 | 1.61 | 1.45 | RMSE | 1.83 | 1.90 | 1.85 | 1.90 |
| Prediction | Pearson's $r$ | 0.55 | 0.65 | 0.61 | 0.65 | Pearson's $r$ | 0.26 | 0.26 | 0.31 | 0.32 |
| Contacts | AUPRC (%) | 23.21 | 23.13 | 25.27 | 24.83 | AUPRC (%) | 12.80 | 10.88 | 9.71 | 9.23 |
| Prediction | AUROC (%) | 82.80 | 82.88 | 82.09 | 82.56 | AUROC (%) | 81.58 | 80.47 | 79.25 | 78.15 |

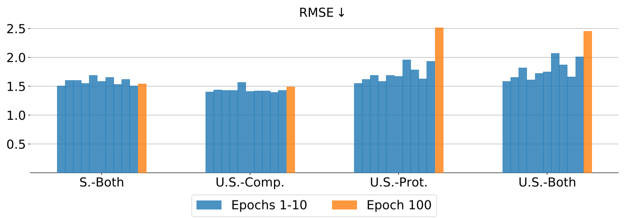


**Fig. S8.** Affinity performance (RMSE) of concatenation model with different epochs pre-trained on the larger set.

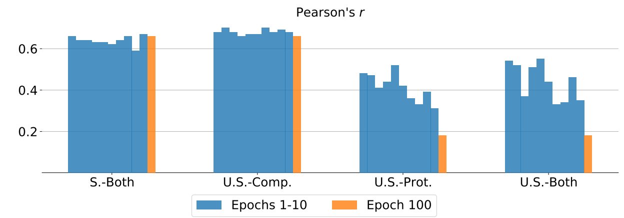


**Fig. S9.** Affinity performance (Pearson's $r$) of concatenation model with different epochs pre-trained on the larger set.

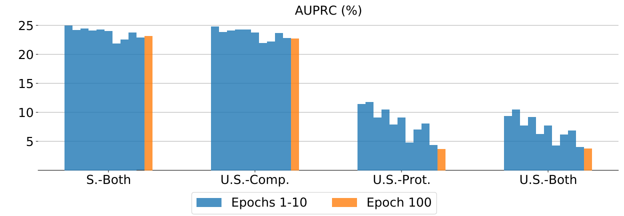


**Fig. S10.** Contact performance (AUPRC) of concatenation model with different epochs pre-trained on the larger set.

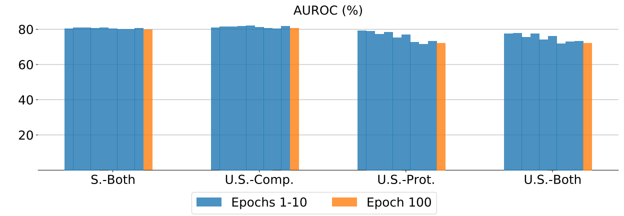


**Fig. S11.** Contact performance (AUROC) of concatenation model with different epochs pre-trained on the larger set.

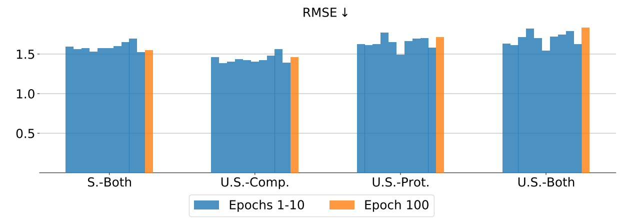


**Fig. S12.** Affinity performance (RMSE) of cross interaction model with different epochs pre-trained on the larger set.

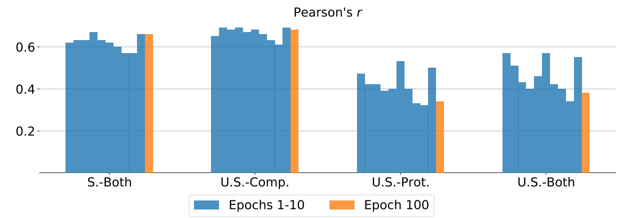


**Fig. S13.** Affinity performance (Pearson's $r$) of cross interaction model with different epochs pre-trained on the larger set.

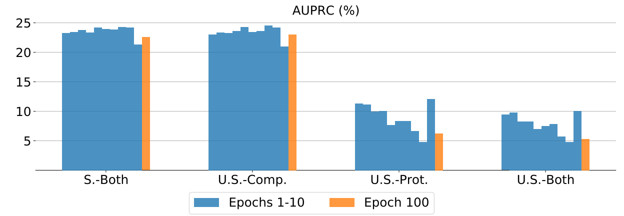


**Fig. S14.** Contact performance (AUPRC) of cross interaction model with different epochs pre-trained on the larger set.

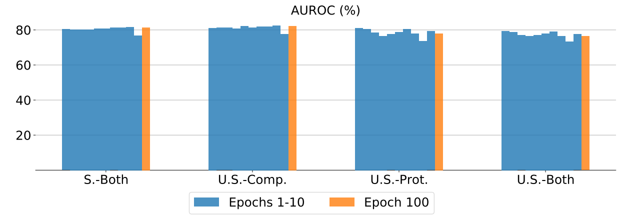


**Fig. S15.** Contact performance (AUROC) of cross interaction model with different epochs pre-trained on the larger set.

optimization settings: the optimizer Adam with a learning rate of 0.0001, the batch size of 32 and the maximum amount of training epochs being 200. The best checkpoint model is selected via validation (we use 10% randomly held-out data of the training set for validation). The following hyperparameters in the loss function are optimized following a two-stage process over pre-defined grids (Karimi *et al.*, 2020). Specifically, $\lambda_{\text{group}}$, $\lambda_{\text{fused}}$, and $\lambda_{\text{L1}}$ are first tuned over {0.01, 0.001, 0.0001} with $\lambda_{\text{cont}} = 0$ (affinity regression alone), where the best affinity loss $l_{\text{aff}}$ is recorded and $\lambda_{\text{group}}$, $\lambda_{\text{fused}}$, and $\lambda_{\text{L1}}$ are optimized with the best AUPRC, such that the corresponding affinity RMSE does not deteriorate more than 10% of the best affinity RMSE. In the second stage, we fix the optimal $\lambda_{\text{group}}$, $\lambda_{\text{fused}}$, and $\lambda_{\text{L1}}$ and tune $\lambda_{\text{cont}}$ over $\{1\text{e}0, 1\text{e}1, 1\text{e}2, 1\text{e}3, 1\text{e}4, 1\text{e}5\}$ based on the best AUPRC performance while jointly optimizing the regularized affinity and contact losses.

When pretraining is additionally introduced, we use two unlabelled datasets at different scales as described in Section 2.1. In the smaller set with ground-truth contact maps for (unbound) proteins, we perform single- and multi-modality pre-training. In the larger set without structure information for proteins, we only pre-train the sequence encoder in the single-modality scheme, and additionally utilize structure information in the smaller set to pre-train the graph encoder in the multi-modality scheme. The masking ratios of sequence and graph self-supervisions are tuned in {0.05, 0.15, 0.25}. The optimization procedure is executed by Adam optimizer with a learning rate of 0.0001 and the batch size is set as 128. We pre-train for 100 epochs in the smaller set, while in the $>$200 times larger set, we observe the "pre-training overfitting" phenomenon based on validation (see Section 3.4) , and thus we cautiously perform pre-training only for one epoch to report the performance. The ablation results on the number of epochs are shown in Appendix F.

### I Procedure of Reproducing MONN

We reproduce MONN (Li *et al.*, 2020) with their public code (https://github.com/lishuya17/MONN) on our curated CPAC dataset. We replace their compound encoder graph warp module (Ishiguro *et al.*, 2019) with a GCN (the same as in DeepAffinity+ (Karimi *et al.*, 2020)) since we treat compound graphs as homogeneous graphs whereas the graph warp module can only be applied on heterogeneous graphs.

The MONN loss is defined as the weighted summation of the affinity and contact losses, i.e. $\text{loss}_{\text{MONN}} = \text{loss}_{\text{aff}} + \alpha\, \text{loss}_{\text{cont}}$ where the weight factor $\alpha$ is pre-set as 0.1. Thereby, we tune hyper-parameters on three configurations as defined in their implementation: (k_head, kernel_size, hidden_size1, hidden_size2) $\in$ {(2, 7, 128, 128), (1, 5, 128, 128), (1, 7, 128, 128)}, with the selection validation criterion: we record the best affinity loss (RMSE), and select the hyper-parameter combination with the best AUPRC such that the corresponding affinity RMSE does not deteriorate more than 10% of the best affinity RMSE.
